# *In situ* reprogramming of gut bacteria by oral delivery

**DOI:** 10.1101/2020.04.15.043232

**Authors:** Bryan B. Hsu, Isaac N. Plant, Lorena Lyon, Frances M. Anastassacos, Jeffrey C. Way, Pamela A. Silver

## Abstract

Abundant links between the gut microbiota and human health indicate that the modification of bacterial function could be a powerful therapeutic strategy. The inaccessibility of the gut and inter-connections between gut bacteria and the host make it difficult to precisely target bacterial functions without disrupting the microbiota and/or host physiology. Herein we describe a multidisciplinary approach to modulate the expression of a specific bacterial gene within the gut by oral administration. We first demonstrate that an engineered temperate phage λ expressing a programmable dCas9 represses a targeted *E. coli* gene in the mammalian gut. To facilitate phage administration while minimizing disruption to host processes, we develop an aqueous-based encapsulation formulation with a microbiota-based release mechanism and show that it facilitates the oral delivery of phage *in vivo*. Finally we combine these technologies and show that bacterial gene expression in the mammalian gut can be precisely modified *in situ* with a single oral dose.

## Introduction

The gut microbiome has numerous associations with human health.^1^ This bacterial community contains hundreds of densely colonizing species with a composition that varies along the gastrointestinal tract, between individuals, and over time.^2^ The complexity of this ecosystem makes it challenging to precisely target specific bacteria without unintended impacts to the microbiota.^3^ To enable the interrogation and therapeutic modification of microbial interactions with the host, generalizable tools are needed, especially ones capable of modifying specific bacterial functions while minimizing disruption to non-targeted genes, microbes, and host physiology.

Multiple biological barriers preclude the efficient modification of bacterial processes within the gut. Oral delivery is the preferable approach but remains challenging because of the acidity and proteases in the upper gastrointestinal tract.^4^ Neutralization of these natural physiological barriers can be disruptive and has been associated with increased secretion due to feedback mechanisms,^5^ susceptibility to enteric pathogens,^6^ and a reduced bacterial diversity in the gut.^7^ Variability in meal timing, intestinal motility and individual-specific physiological conditions can further complicate the efficacy of oral delivery formulations.^8^ Even if these barriers can be overcome, the nature of the microbiota itself, with a dense colonization of competing species, makes the specific and durable modification of bacteria challenging.

Phages are capable of targeting specific bacteria even among a complex consortia. With the prevalence of antibiotic resistant infections, there has been increasing interest in lytic phages for their ability to kill cognate bacteria during phage propagation. Temperate phages, however, have been of less interest therapeutically because they do not primarily pursue lysis and can integrate themselves into the bacterial genomes as prophages. While a potential detriment for bacteriolytic approaches, the specificity and lysogenic conversion of bacteria by temperate phages offers a potential strategy to introduce new genes or even reprogram endogenous gene expression of bacteria within their natural ecosystems. We have previously demonstrated that bacterial virulence can be repressed *in vitro* and in the mammalian gut using a lysogenic phage engineered to specifically repress shigatoxin expression^9^.

Here we report a non-invasive strategy to modify gene expression of specific bacteria in the mammalian gut via oral delivery. First, we engineer temperate phage λ to express a nuclease-deactivated Cas9 (dCas9) that specifically represses gene expression in bacteria both *in vitro* and when colonizing the mouse gut (**Figure 1A**). To improve survival against gastric acid and proteases, program release into the lower GI tract, and minimize potential physiological disruption to host and microbial processes, we develop an aqueous-based encapsulation formulation that protects phage during oral delivery (**Figure 1B**). This biomaterial construct contains a tunable microbiota-specific release mechanism that triggers cargo release upon entry into the bacterial-dense large intestine (**Figure 1C**). Finally, we demonstrate that combining these two technologies enables non-invasive and minimally-disruptive *in situ* modification of bacteria in the mammalian gut.

**Figure 1.**
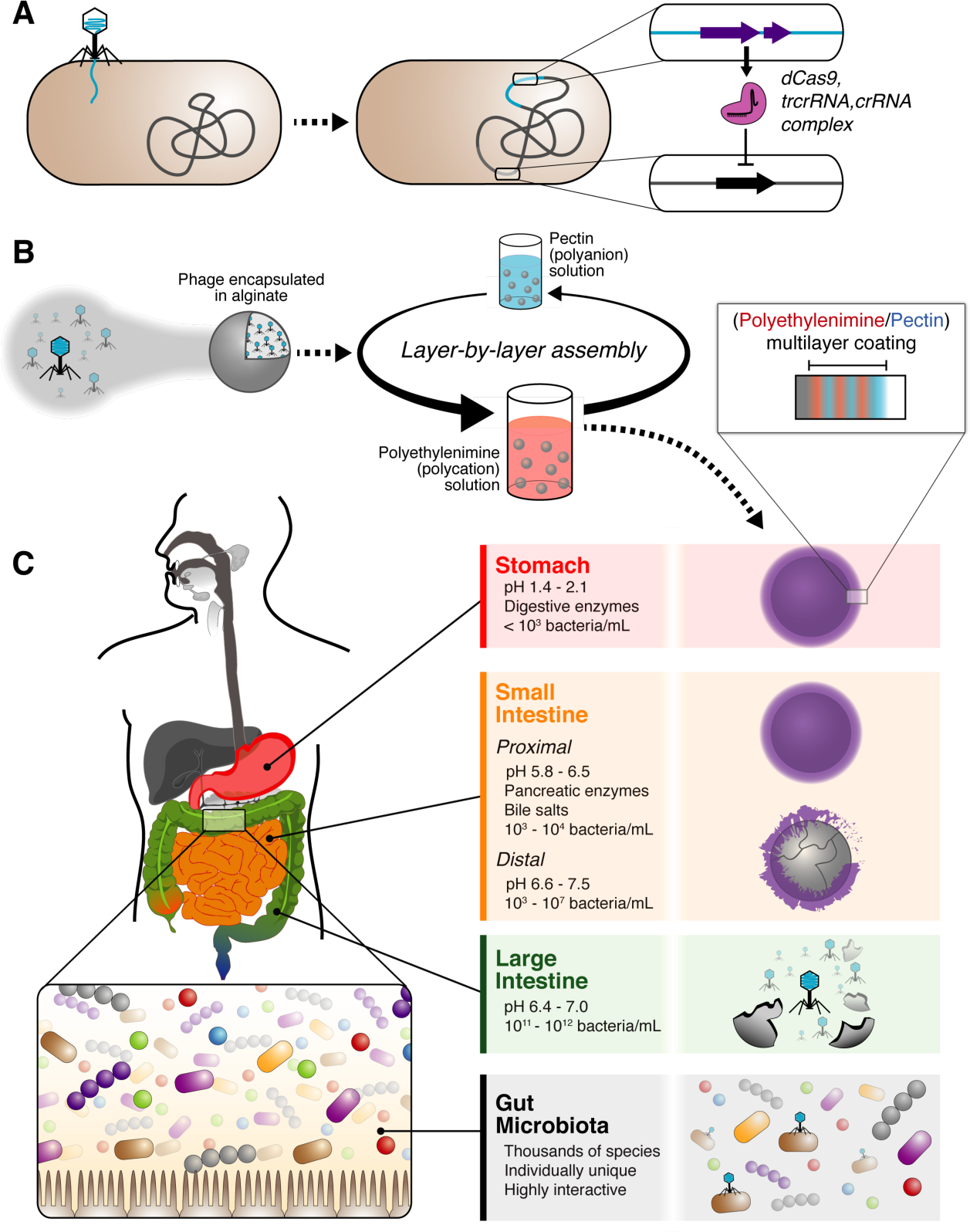
In situ modification of gut bacteria by oral delivery. Temperate phage is engineered to express dCas9, trcrRNA and crRNA so that specific bacterial genes can be repressed (A). The encapsulation strategy uses calcium-mediated crosslinking of a sodium alginate solution to produce beads which are coated in a polymeric multilayer (B). The digestive system has longitudinal variations in pH^10,11^ and bacteria^12^ and presents multiple challenges to biological therapeutics during their transit through the stomach, small intestine and large intestine (C).

## Results and Discussion

### Engineered phage modifies gene expression in vitro

Engineered phage containing dCas9 represses gene function. As shown in **Figure 2A** and **Figure S1**, our engineered construct was inserted into the λ genome by replacing a portion of the non-essential b2 region.^13^ We used chloramphenicol acetyltransferase as the antibiotic resistance and Cas9 from S*taphylococcus aureus*^14^ because of their minimal size. Cas9 was modified with nuclease-inactivating mutations in the HNH and RuvC catalytic regions.^15^ We tested our system in *E. coli* containing genomically-integrated *rfp* and *gfp* genes^16^ and found our plasmid-based *S. aureus* dCas9 was effective in repressing fluorescence (**Figure S2**). When phage containing crRNA targeting *rfp* (λ::dCas9^rfp^) was added to *E. coli* culture, we found that RFP fluorescence was markedly reduced compared to phage lacking this crRNA (λ::dCas9 phage) (**Figure 2B**). *E. coli* cultures receiving buffer showed a transitory decrease in fluorescence from ∼2h to ∼6h compared to those receiving phage, which is due to an absence of initial phage propagation that occurs with the λ::dCas9 and λ::dCas9^rfp^ phage treated samples as confirmed by the bacterial density (**Figure 2C**). Furthermore, the presence of crRNA in λ::dCas9^rfp^ phage does not markedly affect bacterial growth compared to λ::dCas9 phage. To confirm gene repression is maintained once lysogeny is established, we measured RFP fluorescence from lysogens of *E. coli* and confirmed reduced fluorescence in λ::dCas9^rfp^ lysogens compared to both non-lysogens and λ::dCas9 lysogens (**Figure 2D**). Lysogeny did not affect early exponential growth but may affect bacterial density at stationary phase (**Figure 2E**).

**Figure 2.**
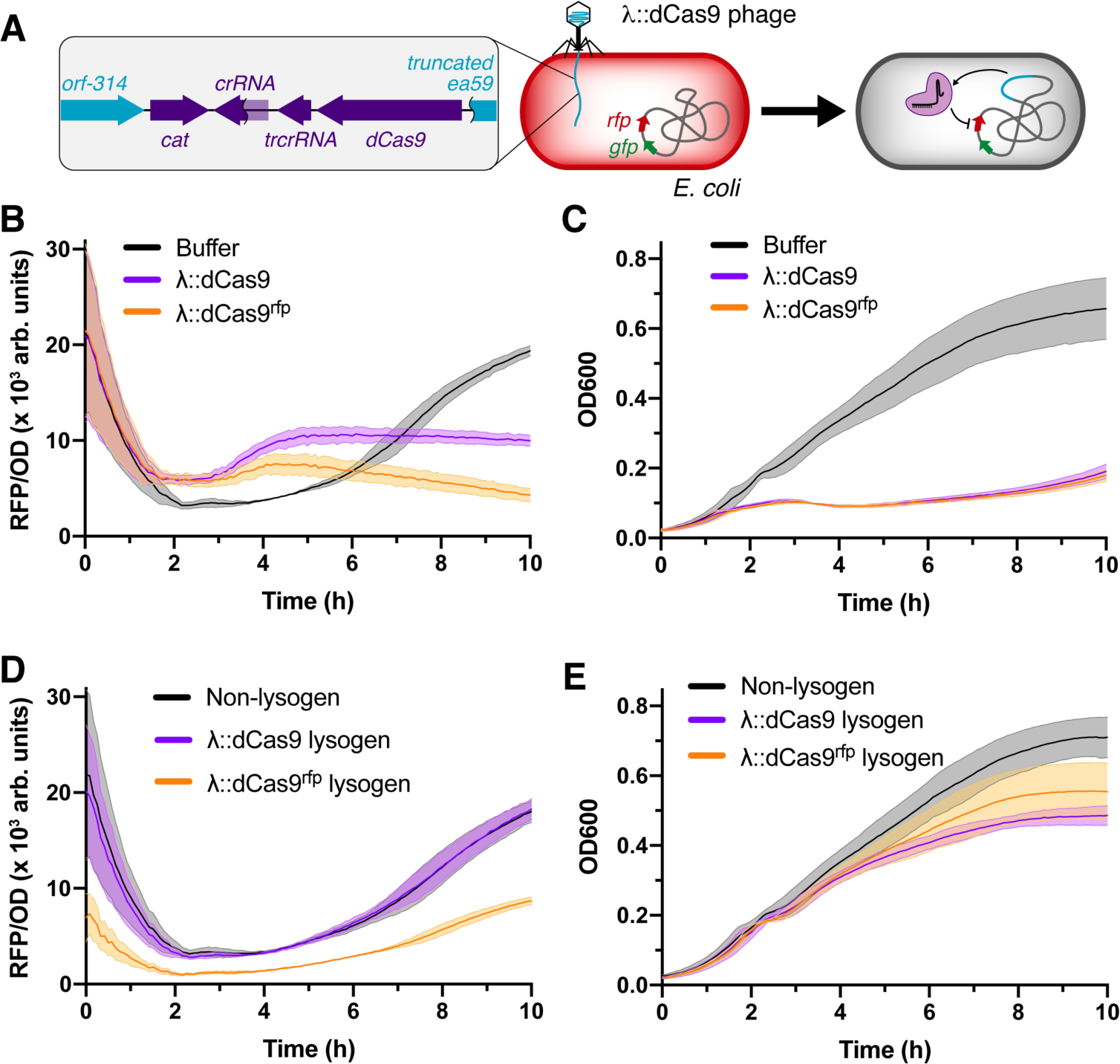
*In vitro* gene repression by λ::dCas9^rfp^. Scheme of *in vitro* experiment examining repression by engineered λ phage (A). *E. coli* cultures mixed with phage buffer, λ::dCas9 phage or λ::dCas9^rfp^ phage tracked for RFP fluorescence (B) and bacterial density (C). Non-lysogenic, λ::dCas9 lysogenic or λ::dCas9^rfp^ lysogenic *E. coli* cultures tracked for RFP fluorescence (D) and bacterial density (E). Lines represent means and shaded regions represent standard errors.

### Engineered phage represses function of gut bacteria in situ

Engineered phage lysogenizes bacteria in the mouse gut. To test the efficacy of our engineered phage *in vivo*, we administered λ::dCas9, λ::dCas9^rfp^ or vehicle (phage buffer) to mice pre-colonized with RFP-expressing *E. coli* and tracked the fecal phage and *E. coli* concentrations over time (**Figure 3A**). To minimize degradation during gastric transit, phage solutions and vehicle were diluted 10-fold into a sodium bicarbonate solution immediately prior to oral gavage. As shown in **Figure 3B**, fecal phage are detectable at high concentrations soon after administration. Despite introduction of phage, total *E. coli* concentrations remain largely consistent (**Figure 3C**) suggesting that a moderate dose (10^7^ pfu) of temperate phage do not markedly affect concentrations of cognate bacteria in the gut. We found that a substantial fraction of fecal *E. coli* are λ::dCas9 or λ::dCas9^rfp^ lysogens soon after phage administration and remain so for the duration of our experiment (**Figure 3D**).

**Figure 3.**
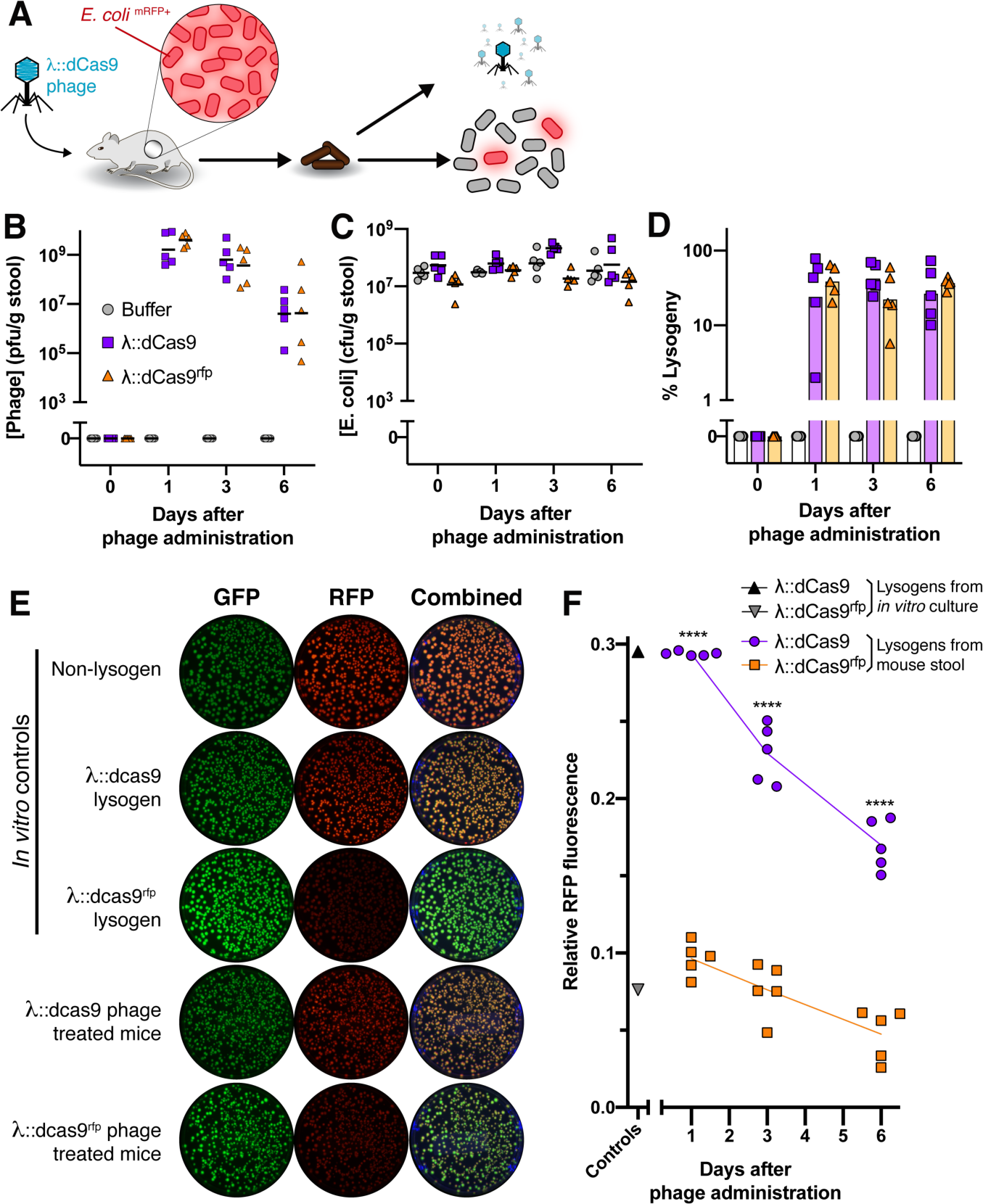
Phage delivered genetic repression *in vivo*. Engineered phage or vehicle was orally administered in bicarbonate solution to mice pre-colonized with *E. coli* expressing RFP and GFP (A). After oral administration, fecal phage (B) total *E. coli* (C) and percentage of *E. coli* lysogenized by phage were quantified (D). Representative fluorescence images of colonies of fecal lysogens are shown compared to *in vitro* cultured controls (E). Relative RFP intensity normalized by GFP intensity of fecal lysogens from mice. Postive and negative controls represent *in vitro* cultured λ::dCas9^rfp^ and λ::dCas9 lysogens, respectively (F). Symbols represent individual mice (*n* = 5) with bars or lines indicating the median. Significance was calculated for preselected time- matched samples using one-way ANOVA with a Sidak post hoc test (****, P < 0.0001).

Lysogenized *E. coli* have reduced fluorescence. We assessed functional gene repression by our engineered phage by measuring the relative RFP fluorescence of λ::dCas9 and λ::dCas9^rfp^ lysogens isolated from mouse stool after oral phage administration. As shown in the representative culture plates in **Figure 3E**, fecal *E. coli* colonies from mice receiving λ::dCas9^rfp^ phage demonstrate a maintained GFP and reduced RFP fluorescence compared to fecal colonies from mice receiving λ::dCas9 phage. These features were similar to *in vitro* cultured λ::dCas9^rfp^ or λ::dCas9 lysogens, respectively. An ensemble view of the relative fluorescence of ∼50 fecal colonies from each mouse at each timepoint shows that RFP repression is maintained in each mouse over time (**Figure S3**) with significant fluorescence reduction by λ::dCas9^rfp^ phage compared to λ::dCas9 phage (**Figure 3F**).

### Thin film coatings resist acid penetration

To minimize degradation during oral delivery in the absence of buffering agents, we developed an aqueous-based encapsulation formulation capable of resisting acid penetration. Alginate beads were generated by adding an alginate solution dropwise to a stirring calcium chloride solution, which mediates gelation through ionic crosslinking. With coaxial air flow we could tune bead size **Figure S4**. Using a Layer-by-Layer (LbL) assembly technique in which polyelectrolyte multilayer films are built by the alternating deposition of polycations and polyanions from aqueous solution (**Figure 1B**), we coated these beads with polyethylenimine (PEI) and pectin to form thin films denoted (PEI/pectin)_n_ where *n* represents the number of “bilayers” deposited. On a flat substrate, these films show the characteristic exponential growth profile for weak polyelectrolytes in high salt solutions (**Figure S4**).^17^ Pectin is included as bacteria-specific degradation mechanism to trigger bead degradation once reaching the gut. We confirmed that CaCl_2_, which is needed to ensure phage and alginate bead stability, did not disrupt polyelectrolyte complexation (**Figure S5**), and that these films remained intact in the presence of simulated gastric fluid (SGF) with pepsin (**Figure S6**).

LbL-film coated alginate beads are resistant to external pH changes. We quantified the acid resistance of these thin films by encapsulating an Oregon Green 488—dextran conjugate, which contains a pH sensitive dye that fluoresces under neutral pH and quenches in acid (**Figure S7**). When suspended in SGF, pH 1.1, the fluorescence quenching of these beads indicates the acidification of the bead interior from its initially physiological conditions of pH 7.5 (**Figure 4A)**. As shown in **Figure 4B**, acid readily infiltrates the entirety of uncoated alginate beads (0-BL) within ∼30 seconds. Increasing the number of bilayers, *i.e*. film thickness, increases acid resistance. Quantification of relative fluorescence intensity reveals that increasing the number of bilayers deposited increases acid resistance with 15 and 15.5-BL coatings maintaining nearly unchanged internal conditions during the first 90s (**Figure 4C**). Furthermore, the acid resistance is retained during prolonged incubation of 2 hours at 37°C (**Figure S8**). Encapsulating λ phage into alginate beads coated with (PEI/pectin)_15.5_ films revealed superior *in vitro* stability against acid compared to λ phage in uncoated alginate beads and non-encapsulated free λ phage (**Figure S9**).

**Figure 4.**
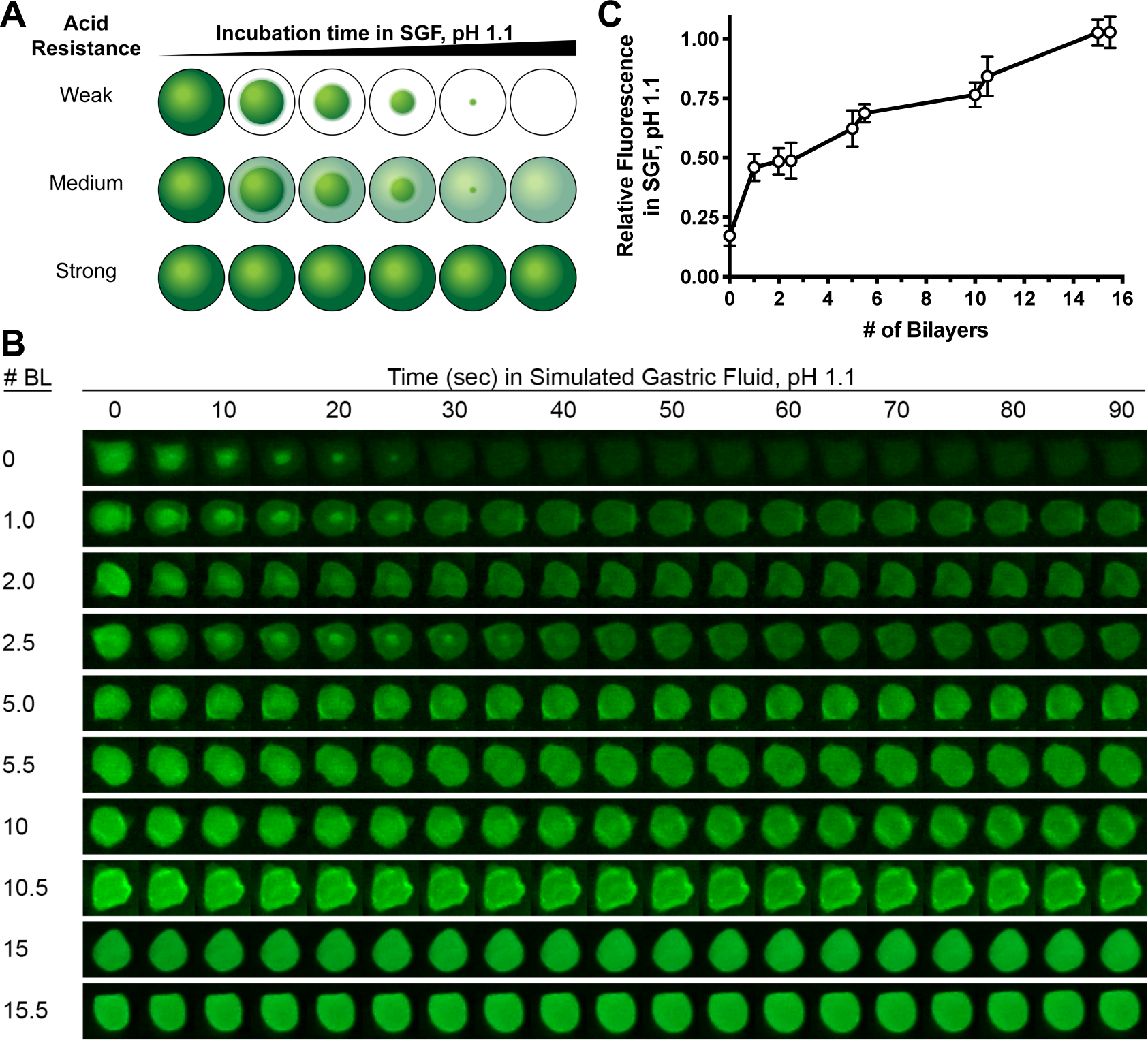
Acid resistance of alginate beads. Encapsulation of a pH sensitive dye in the beads can gauge the pH within the beads after incubation in simulated gastric fluid (SGF) by a large or small loss in fluorescence indicating beads that have been constructed with weak, medium, or strong acid resistance (A). By tracking the fluorescence behavior of representative beads, non-coated (0-BL) and coated (up to 15.5-BL) alginate beads show distinct differences in acid resistance (B). The normalized change in fluorescence after incubation in SGF, pH 1.1 for 90 s shows a relationship with the number of bilayers deposited (mean ± s.d.) (C).

### Encapsulation protects phage during oral delivery

Non-encapsulated free phage is readily degraded when administered orally to mice. For a baseline measure of the degradative conditions of the stomach, we administered increasing concentrations of λBH1 phage in water to mice pre-colonized with *E. coli* MG1655. λBH1 phage is an engineered λ phage encoding an antibiotic resistance marker allowing us to quantify lysogens by antibiotic selection.^9^ Overnight fasting can raise the gastric pH (pH ∼4)^18^, so to preserve gastric acidity, we gavaged mice in the fed state to keep the gastric conditions (pH ∼3)^18^ which is closer to that of humans (pH ∼2).^10^ We found the mouse gastric pH to be 2.39 ± 0.56 (mean ± s.d., n = 5). After administering steadily increasing doses (10^1^ pfu to 10^8^ pfu) of λ phage suspended in water, we monitored fecal phage and lysogen content in mouse stool for 3 days. As shown in **Figure S11**, no phages or lysogens were detected in the feces for doses up to 10^7^ pfu, despite the persistent colonization by *E. coli* that would readily amplify any phage surviving oral administration. It is not until a dose of 10^8^ pfu that phage and lysogens are detected in the stool. The necessity of such a high dose indicates λ phage is readily degraded when orally administered *in vivo*.^19^

LbL-coated encapsulation formulations protect phage during orally delivery in mice. To determine the protective efficacy of our encapsulation formulation during oral administration, mice were gavaged with a low dose (10^3^ pfu) of phage in various formulations suspended in water (**Figure 5A**). Due to size restrictions in mouse anatomy, we used 0.65 mm diameter alginate beads (**Figure S4**). As shown in **Figure 5B**, oral administration of phage encapsulated in alginate beads coated with (PEI/pectin)_15.5_ films resulted in phage detected in stool for all mice within 2 d and continued so for 2 wks (**Figure S12**). By contrast, phage alone or phage co-administered with empty alginate beads coated with (PEI/pectin)_15.5_ films did not survive oral administration (**Figure 5A**). The latter indicates that encapsulation is necessary, and that the beads do not provide protection by somehow inactivating the digestive process. Similarly, phage encapsulated in alginate beads without the LbL coating were completely inactivated, confirming that the LbL films are the essential protective element.

**Figure 5.**
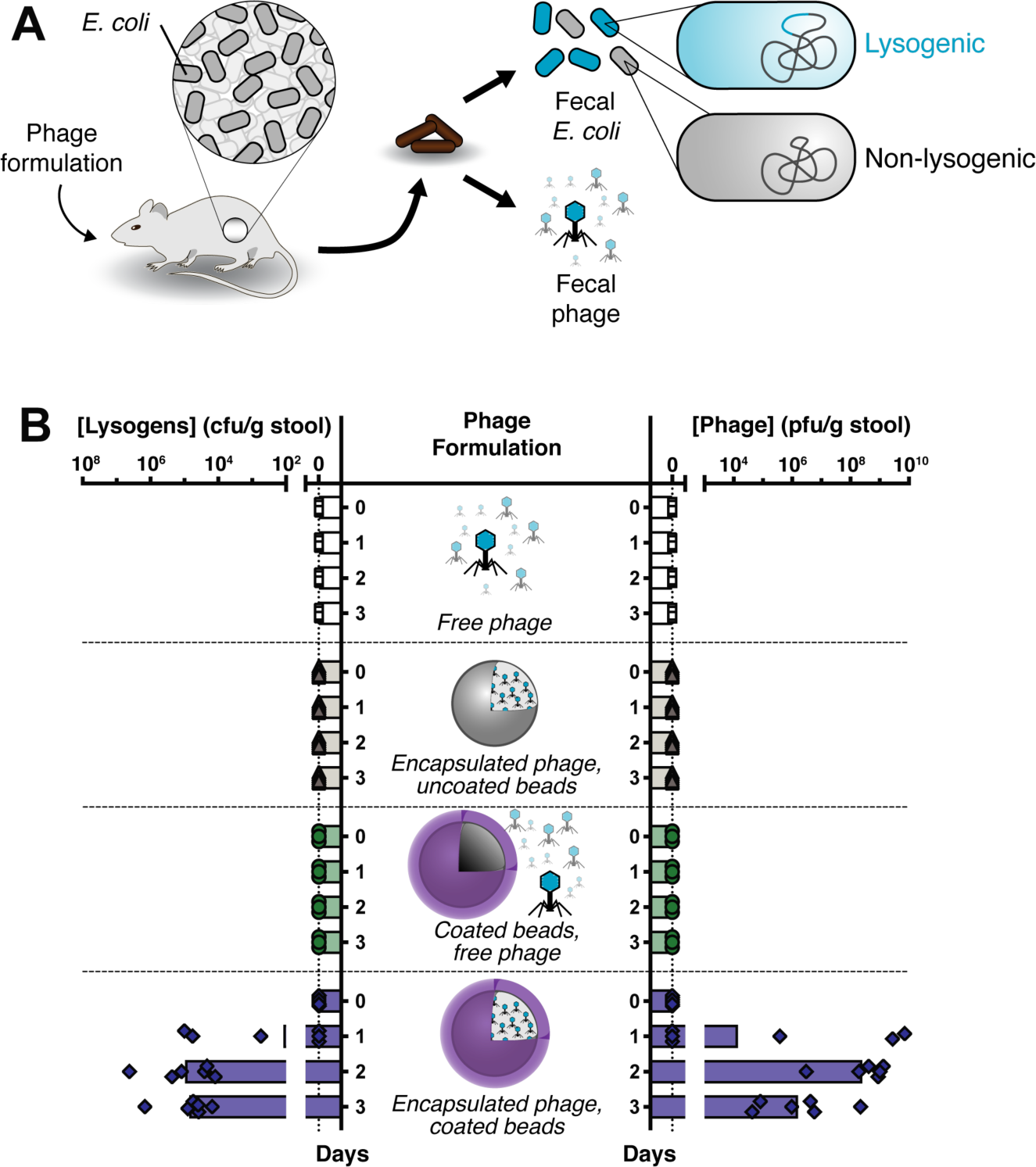
Efficacy of oral encapsulation formulation on bacteriophage survival in mice. Mice pre-colonized with *E. coli* were orally administered phage formulations (A). Mice were administered either free phage (n=5), phage encapsulated in uncoated beads (n=6), a mixture of free phage with empty LbL-coated beads (n=3), or phage encapsulated in LbL-coated beads (n=6). “LbL-coated” indicates coatings of (PEI/pectin)_15.5_ films. Fecal concentrations of lysogens and phage were quantified over time (B). Symbols represent individual mice with bars indicating median.

### Non-invasive and minimally disruptive modification of gut bacteria

Encapsulated phage modifies gut bacteria *in situ* without compromising the gastric barrier. In **Figure 3**, we demonstrate that λ::dCas9^rfp^ phage significantly represses RFP fluorescence in gut bacteria compared to λ::dCas9 phage, which lacks targeting to the *rfp* gene of *E. coli*. In **Figures S11 and 5**, we demonstrate that phage administered in water is readily inactivated during oral administration unless the stomach acid is neutralized, phage is given at very high concentration, or phage is encapsulated into our LbL-coated alginate beads. To determine whether encapsulation impairs λ::dCas9^rfp^ phage performance *in situ*, we administered a low concentration (5 x10^3^ pfu) of this phage either in LbL-coated beads suspended in water or as free phage mixed with bicarbonate buffer to neutralize the stomach acid (**Figure 6A**). Examination of mouse stool after oral phage administration reveals similar fecal phage concentration (**Figure 6B**), total fecal *E. coli* concentration (**Figure 6C**), and lysogenic conversion of fecal *E. coli* (**Figure 6D**). When examining the relative RFP fluorescence of individual λ::dCas9^rfp^ lysogen colonies, buffered and encapsulated phage both provided similar levels of RFP repression for the duration of the study (**Figure 6E**).

**Figure 6.**
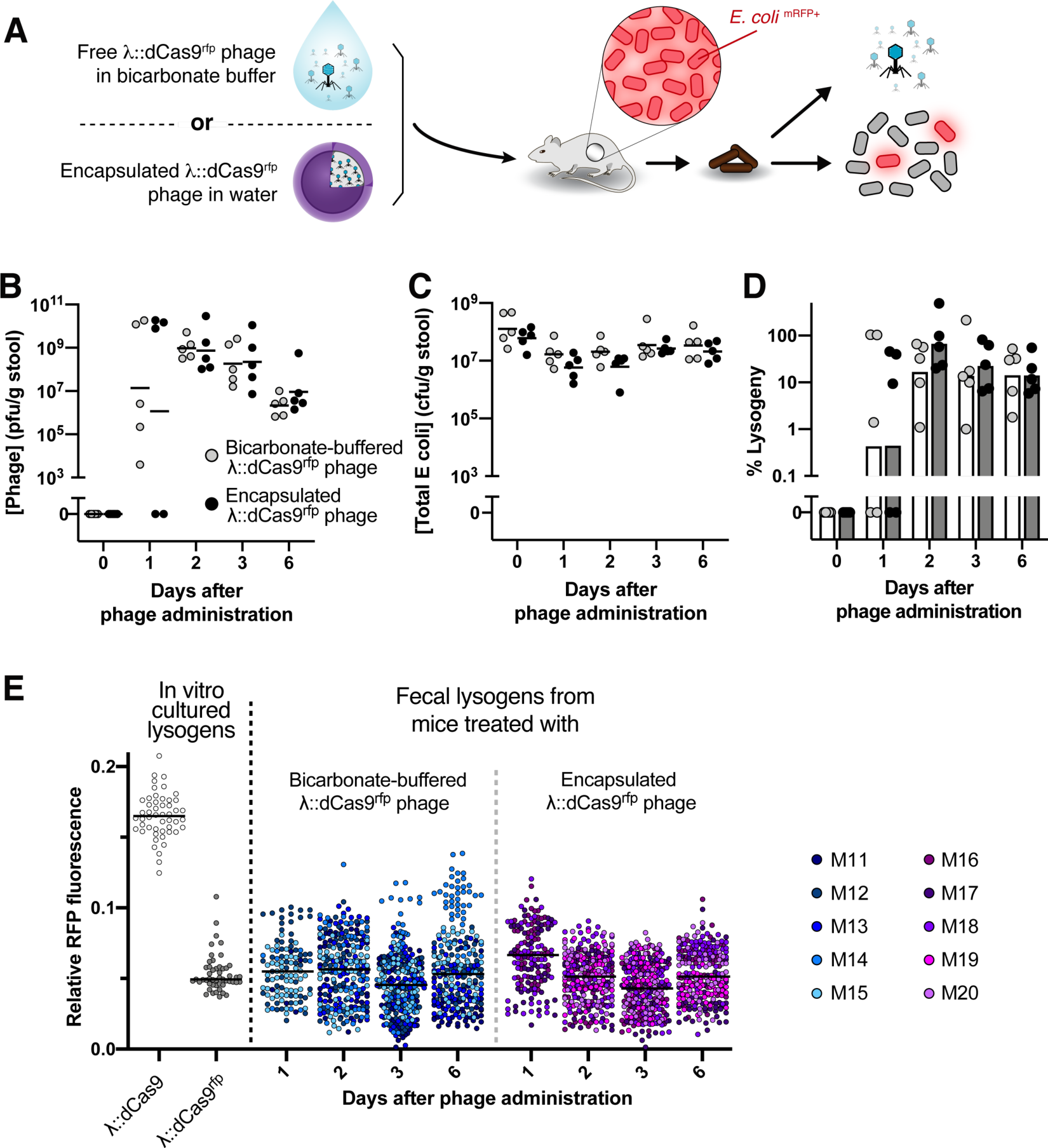
Encapsulation protects oral phage but does not impair function. Free λ::dCas9^rfp^ phage in bicarbonate buffer (to ensure survival) or encapsulated λ::dCas9^rfp^ phage in water were orally administered to mice pre-colonized with *E. coli* expressing RFP and GFP (A). After oral administration, fecal phage (B) total E. coli (C) and percentage of *E. coli* lysogenized by phage were quantified (D). Symbols represent individual mice (*n* = 5) with bars or lines indicating the median. Ensemble view of relative fluorescence from lysogens shown with *in vitro* cultured controls (E). Symbols represent individual colonies (∼50 colonies per mouse sample or culture).

## Discussion

Herein we demonstrate microbiological and biomaterial advances that, while versatile on their own, can be combined to maximize precision targeting of bacteria in the gut. Using temperate phage and RNA-guided dCas9, we describe a precise and programmable approach for the *in-situ* modification of bacteria and bacterial transcription. To maximize the efficacy of oral delivery while minimizing potential physiological disruption, we developed an acid-resistant encapsulation formulation that relies on endogenous bacterial enzymes in the lower GI tract to trigger release of engineered phage. Combined, these technologies demonstrate a non-invasive, durable, and targeted approach towards modifying bacterial function in the gut.

Manipulating bacterial processes *in situ* could be a powerful tool for characterizing the role of specific bacterial genes in host health. Links between microbial metabolites and human disease have spawned interest in compounds that specifically target bacterial functions^20^ even though the compounds have potential off-target effects^21^. As a research tool, our approach offers a potential strategy to systematically screen putative targets, characterize cause-effect relationships, and identify cascading effects to the surrounding microbiota and host. As a therapeutic, our work outlines a generalizable strategy for durable modification of gut bacteria using a single, self-propagating dose.

With this exquisite specificity comes a possibly narrow efficacy. Since microbial functions can be shared across co-colonizing bacteria it may be necessary to modify more than one species to achieve systemic changes. Analogous to traditional phage therapy, cocktails could be used to broaden the host range of targeted bacteria. Another general consideration when deploying phages is the development of resistance. Prophages are prevalent among gut bacteria ^22^ and likely persist because they confer fitness advantages to the bacterial host^23^ suggesting that engineered temperate phages could similarly be tolerated in the gut microbiota.

As we gain greater insight into the extent of host-microbiome interactions, the development of precision tools to modify the function of species within their ecosystem could improve both the specificity and durability of the intervention. While there is great interest in the human gut, the microbiomes of other bodily regions such as the mouth, skin, and lungs similarly have therapeutic potential. In addition to other hosts such as mammals, plants, and insects, the microbiota of soil and marine environments could also benefit from the modification of specific microbes. With the tool we have described here, such modifications have become more realistic.

## Acknowledgements

We are grateful to Dr. Georg Gerber for insightful discussions and Amanda Graveline, DVM for her assistance and training for mouse experiments. We are also grateful to Calixto Saenz and the HMS Microfabrication Core Facility for assistance with the microfluidic devices. This work was supported by Defense Advanced Research Projects Agency Grant HR0011-15-C-0094 and funds from the Wyss Institute for Biologically Inspired Engineering. B.B.H. is grateful for support from the Rosenbloom Postdoctoral Fellowship.

## Author Contributions

Conceptualization of engineered phage, B.B.H. and I.N.P.; Conceptualization of encapsulation formulation, B.B.H.; Visualization, B.B.H.; Methodology, B.B.H. and I.N.P.; Investigation, B.B.H., I.N.P., L.L., and F.A.; Funding Acquisition, J.C.W. and P.A.S.; Writing – Original Draft, B.B.H.; Writing – Review and Editing, I.N.P. and P.A.S.

## Materials and Methods

### Molecular cloning and phage engineering

Golden Gate Assembly was used for cloning plasmids (**Figure S1**). Q5 Hot Start polymerase was used to amplify the proC promoter, λ phage homology arms, pACYCDuet origin of replication, and pACYCDuet chloramphenicol resistance cassette. Restriction sites were also added during PCR. A DNA sequence including the coding sequence of *S. aureus* dCas9, the dCas9’s tracRNA under a constitutive promoter, and the dCas9’s crRNA under a constitutive promoter, was ordered from IDT as two gBlocks. Golden Gate reactions were run using 10xT4 Ligase Buffer (Promega), T4 Ligase (2,000,000 units/mL, NEB), and BSA (10 mg/mL, NEB) as well as the appropriate restriction enzyme, either Eco31I, Esp3I, or SapI (Thermo FastDigest). Golden Gate reactions were desalinated using drop dialysis (for a minimum of 10 minutes) and electroporated in DH10β Electrocompetent Cells (Thermo Fischer). Plasmids were verified by sequencing all junctions as well as the entirety of the gBlocks. The gRNA spacers were synthesized as complementary oligos then added to plasmids by Golden Gate after being annealed and phosphorylated. This was done by incubating the oligos with T4 Ligase Buffer (NEB) and T4 Poly Nucleotide Kinase (NEB), heating to boiling, and then slowly cooled to room temperature. Annealed oligos were added to plasmids using Eco31I. The gRNA spacers were verified by sequencing.

To generate a crude phage lysate containing the desired recombinant phage, cultures of log-phase *E. coli* C600 containing dCas9 plasmids grown in TNT (tryptone-NaCl-thiamine) media with 25 ug/mL chloramphenicol were pelleted and resuspended into an equal volume of TNT media. Plasmids contained 40-bp homology regions to *ea59* and *orf-314* genes to facilitate homologous recombination of the dCas9 construct into λ phage. In a double agar layer plaque assay method^26^, 100 µL of this culture was mixed with 100 µL of serially diluted λ phage in phage buffer (50 mM tris, 100 mM sodium chloride, 10 mM magnesium chloride, and 0.01% gelatin at pH 7.5) and then mixed with 3 mL of molten top agar (TNT with 0.3% agar) and immediately poured onto TNT agar plates to harden. After overnight culture at 37°C, the top agar from plates containing the highest density of individual plaques were resuspended into 5mL of phage buffer with gentle rocking at 4°C for 2h. These suspensions were then pelleted with the supernatant filtered through 0.45 um syringe filters, yielding crude phage lysates.

*E. coli* C600 grown to late log in TNT with 0.4% maltose was concentrated by pelleting and resuspension to ∼10^10^ cfu/mL. 200 µL of this bacterial suspension was mixed with 200 µL of crude phage lysate and incubated at 37°C for 2.5 h, statically. Cultures were then plated onto LB with 34 ug/mL chloramphenicol and incubated overnight at 37°C. For plaque purification, phage was isolated by culturing streak-purified colonies in TNT overnight at 37°C, then treated with chloroform and pelleted. Plaques were then generated from serial dilutions of the supernatant by double overlay plaque assay with *E. coli* (*rfp*^+^, *gfp*^+^). After overnight incubation at 37°C, plaque centers were picked and streaked onto LB with chloramphenicol. Resultant colonies were checked for GFP fluorescence and PCR amplicons to confirm the correct bacterial host and presence of phage, respectively.

### In vitro fluorescence measurements

In flat bottom 96-well fluorescence plates, 180 uL of log-phase *E. coli* (*rfp*^+^, *gfp*^+^) cultured in TNT and diluted to OD600 ∼ 0.05 (1-cm pathlength) were mixed with 20 µL of λ::dCas9 phage or λ::dCas9^rfp^ phage for a final MOI∼1.0, or phage buffer. Plates were shaken at 37°C for 10 h with measurements of OD_600_, GFP fluorescence (ex 485nm/em 528nm), and RFP fluorescence (ex 555nm/em 584 nm) at 5 min intervals (BioTek Synergy H1MF). Studies of non-lysogens, λ::dCas9 lyosgens or λ::dCas9^rfp^ lysogens were setup similarly except that 200 µL of log-phase cultures were used.

### Bead generation and film coating

Alginate beads were generated in a similar manner as described previously^24^. A 1% sodium alginate solution in phage buffer was dissolved with stirring, heated until boiling, and then cooled to 4°C. This solution was added drop-wise with a 30 gauge needle (BD) at 50 mL/hr via syringe pump (New Era Pump Systems) to a stirring solution of phage buffer containing 100 mM calcium chloride. After complete addition, the bead suspension was stirred for an additional hour, then washed twice with phage buffer containing 100 mM calcium chloride and stored in the same solution at 4°C. For encapsulation, bacteriophage was included in the sodium alginate solution prior to addition to the calcium chloride bath. Average bead diameters were determined from at least one hundred beads using ImageJ.

We then coated these alginate beads in polyelectrolyte multilayer films. For one bilayer, beads were incubated for 15 min in a polycation solution and washed twice, then incubated for 15 min in a polyanion solution and washed twice. This process was repeated *n*-times for *n*-bilayers. The polycation solution consisted of 2 mg/mL branched polyethylenimine (PEI, 750 kDa) and the polyanion solution consisted of 2 mg/mL apple pectin. Both polyelectrolyte solutions and wash buffer were formulated in phage buffer with 100 mM calcium chloride, pH 7.5. After film deposition, beads were stored at 4°C.

### Film characterization

Multilayer films were assembled onto silicon wafers (University Wafer) using microfluidic devices fabricated using soft-lithography techniques as described previously.^25^ The photolithographic mask used is shown in **Figure S13**. Multilayer films were assembled in devices by adding 10 µL of 2 mg/mL PEI solution in phage buffer for 5 min, washing twice with phage buffer, then adding 10 µL of 2 mg/mL pectin solution in phage buffer for 5 min, and washing twice with phage buffer. At the end of step, solutions were removed from wells by aspiration. Film thicknesses were measured by profilometry (KLA-Tencor).

### In vitro characterization of bead function

We assessed the internal pH of the alginate beads by incorporation of 150 µg/mL (∼100 µM dye) of the polymer-dye conjugate, dextran-Oregon Green 488 (70kDa M_W_, Life Technologies), in the sodium alginate solution prior to gelation. In 24-well plates, we re-suspended the beads into 1 mL of simulated gastric fluid (SGF), pH 1.1 and imaged the fluorescence using a macroscope device, as described previously^26^. SGF consisted of 34 mM sodium chloride and 85 mM HCl, pH 1.1. The relative change in fluorescence for each bead was determined by first subtracting the raw background intensity immediately adjacent to the beads from the raw intensity within the center of the beads and then taking the relative change in fluorescence as determined by the fraction of remaining fluorescence after 90 s compared to its initial fluorescence, immediately after the addition of SGF. For the qualitative comparison in change of fluorescence over time, identically adjusted the contrast and brightness for each bead montage over time was used.

We determined the in vitro protective ability of encapsulation by suspending individual beads in 0.5 mL of SGF (pH ∼1.1 to 7.4) for 15 min at 37°C. Bacteriophage was released by re-suspending the beads into phage buffer containing 0.5 mg/mL alginate lyase and 5 mg/mL pectin lyase with shaking at 37°C for 10 min with mechanically disruption using a sterile wooden stick, and then continued shaking at 37°C for an additional 10 min. This process did not affect phage titers (**Figure S10**) Free bacteriophage was diluted 1000-fold into 0.5 mL of SGF (pH ∼1.1 to 7.4), incubated for 15 min at 37°C, and then diluted 50-fold into phage buffer. Samples were kept on ice until used in for the plaque assays described above.

### Animal studies

All animal work was approved by the Harvard Medical School IACUC under protocol 04966. Upon arrival, 6-7 week old female BALB/c mice (Charles River) were acclimated for a week prior to experiments. Similar to as previously described, 5 g/L streptomycin sulfate USP grade (Goldbio) was provided in the drinking water one day prior to oral gavage with 100 µL of streptomycin resistant *E. coli* MG1655 or *E. coli* (*rfp*^+^, *gfp*^+^) using 20 gauge PTFE animal feeding needles (Cadence Science). The gavage solution of E. coli was prepared by inoculating an overnight culture (∼16-20 h) in LB with 100 µg/mL streptomycin, then washing twice by pelleting and then re-suspending in an equal volume of PBS, and then diluting 10-fold into PBS to yield ∼10^7^ – 10^8^ cfu/mL. One day after administration of E. coli, 100 µL of phage solution was administered by oral gavage.

Testing the efficacy of engineered phage for repressing RFP fluorescence (**Figure 3**). Solutions of vehicle (phage buffer), either 10^9^ pfu/mL of λ::dCas9 phage or 10^9^ pfu/mL of λ::dCas9^rfp^ phage, were diluted 10-fold into 0.1 M sodium bicarbonate followed by immediate administration of 100 µL to mice by oral gavage.

Testing the susceptibility of phage to inactivation during oral delivery (**Figure S11 and Figure 5**). To test free phage, λBH1 was diluted into water and then 100 µL was administered to mice by oral gavage. Increasing doses of 10^1^, 10^3^, 10^5^, 10^7^, and 10^8^ pfu were administered at three-day intervals. λBH1 phage or lysogens were not detected in stool until after the highest dose. To test susceptibility of phage in different formulations, λBH1 phage free in solution, mixed (i.e. not encapsulated) with LbL-coated alginate beads, encapsulated into alginate beads without coating or encapsulated into LbL-coated alginate beads. The LbL coating consisted of a (PEI/pectin)_15.5_ film. Phage solutions were diluted into water immediately prior to oral administration of 100 µL to mice while encapsulated phage was resuspended into water immediately prior to oral administration of 100 µL containing 50 beads. All solutions and suspensions were administered using 16 gauge polyurethane feeding needles (Instech Labs) to enable administration of beads.

Testing the efficacy of encapsulated engineered phage in repressing RFP fluorescence (**Figure 6**). To test the efficacy of encapsulated λ::dCas9^rfp^ phage, 100 µL of a low-dose (5 ×10^3^ pfu) of free phage or phage encapsulated into LbL-coated beads with (PEI/Pectin)_15.5_ films were administered to mice by oral gavage. Free λ::dCas9^rfp^ phage was diluted 10-fold into 0.1 M sodium bicarbonate immediately before administration to mice. Encapsulated phage (in 7 beads) was resuspended into water immediately before administration to mice. Solutions and suspensions were administered by oral gavage using 16 gauge polyurethane feeding needles (Instech Labs) to enable administration of beads.

To quantify the colonization within the mouse gut, daily stool samples were obtained. To quantify phage, within 30 min of excretion stool was suspended into 1 mL of phage buffer using sterile hospital sticks and then kept on ice. After 10 min, a few drops of chloroform was added to kill bacteria without affecting the bacteriophage.^26^ After an additional 10 min, the solution was pelleted at 4,000 rpm for 10 min and then the supernatant diluted into phage buffer and quantified by plaque assay against the indicator bacteria, *E. coli* C600 using a double agar layer approach. To quantify bacteria, stool was frozen at -80°C within 30 min of excretion and then immediately prior to analysis, thawed at room temperature, re-suspended into 1 mL of PBS with vortexing for 10 min at 4°C and then gently centrifuged at 200 rpm for 20 min to allow debris to settle while leaving bacteria in suspension. E. coli was quantified by plating 100uL of serial 10-fold dilutions in PBS onto MacConkey agar (Remel) supplemented with (100 μg/mL) streptomycin to quantify total E. coli or MacConkey agar supplemented with (100 μg/mL) streptomycin sulfate and (50 μg/mL) kanamycin sulfate to quantify bacteriophage lysogens. Fluorescence from *E. coli* colonies was measured by plating onto LB with 34ug/mL chloramphenicol and incubated at 37°C for 2 d. Fluorescence images were taken with a Bio-Rad alpha imager and fluorescence intensity quantified by ImageJ.

To determine the gastric pH, mice were allowed free access to food and water, then sacrificed under CO2 and cervical dislocation, dissected, and a pH probe was immediately inserted into the gastric contents for measurement.

## Supplementary Information

**Figure S1.**
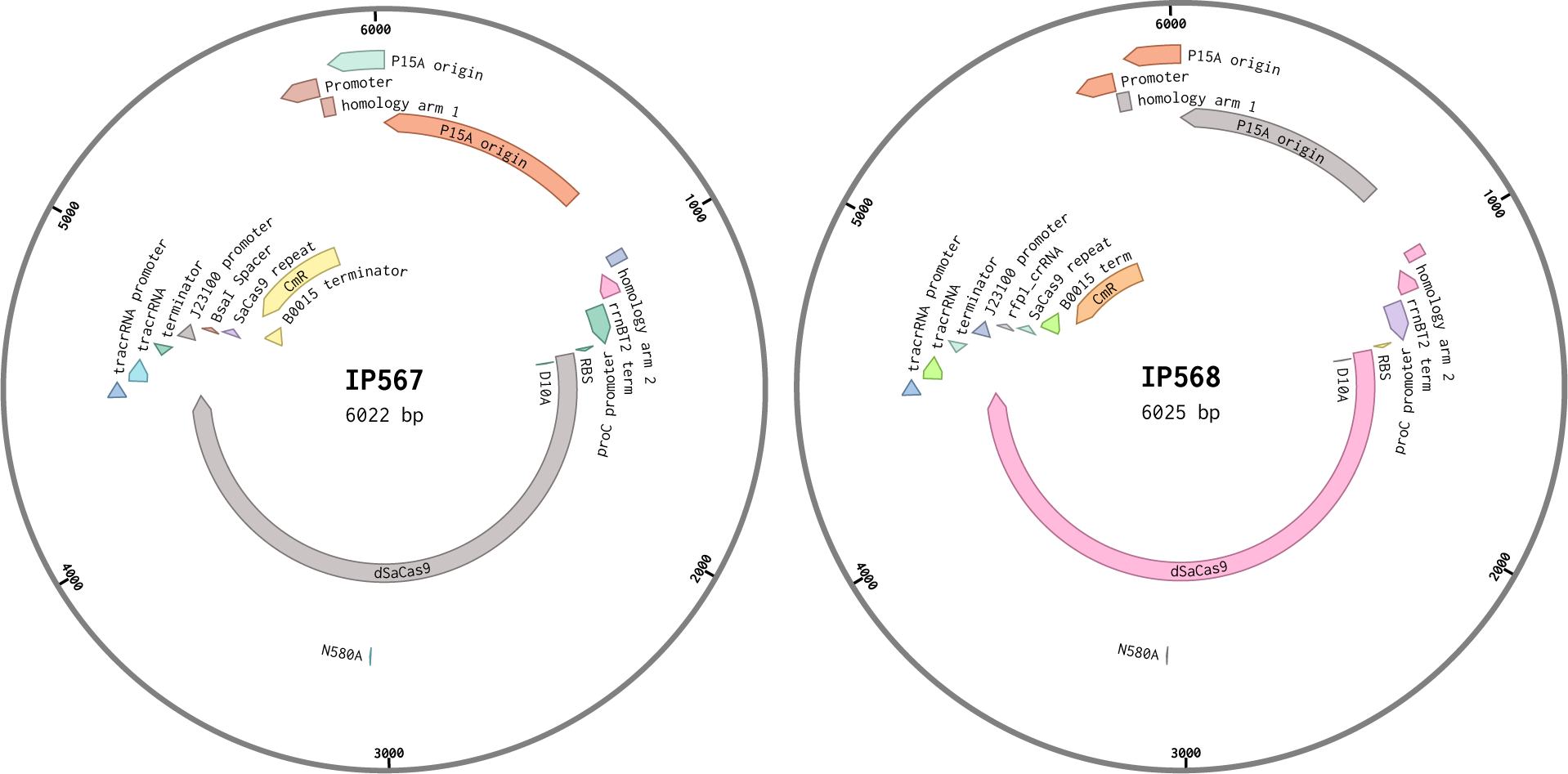
Plasmid maps. Plasmids containing dCas9, tracrRNA and chloramphenicol resistance constructs without (IP567) and with crRNA targeting *rfp* (IP568) were cloned into a pACYCDuet backbone. Homology arms 1 and 2 corresponded to regions in *orf-194/orf-314* and *ea59* in λ phage, respectively.

**Figure S2.**
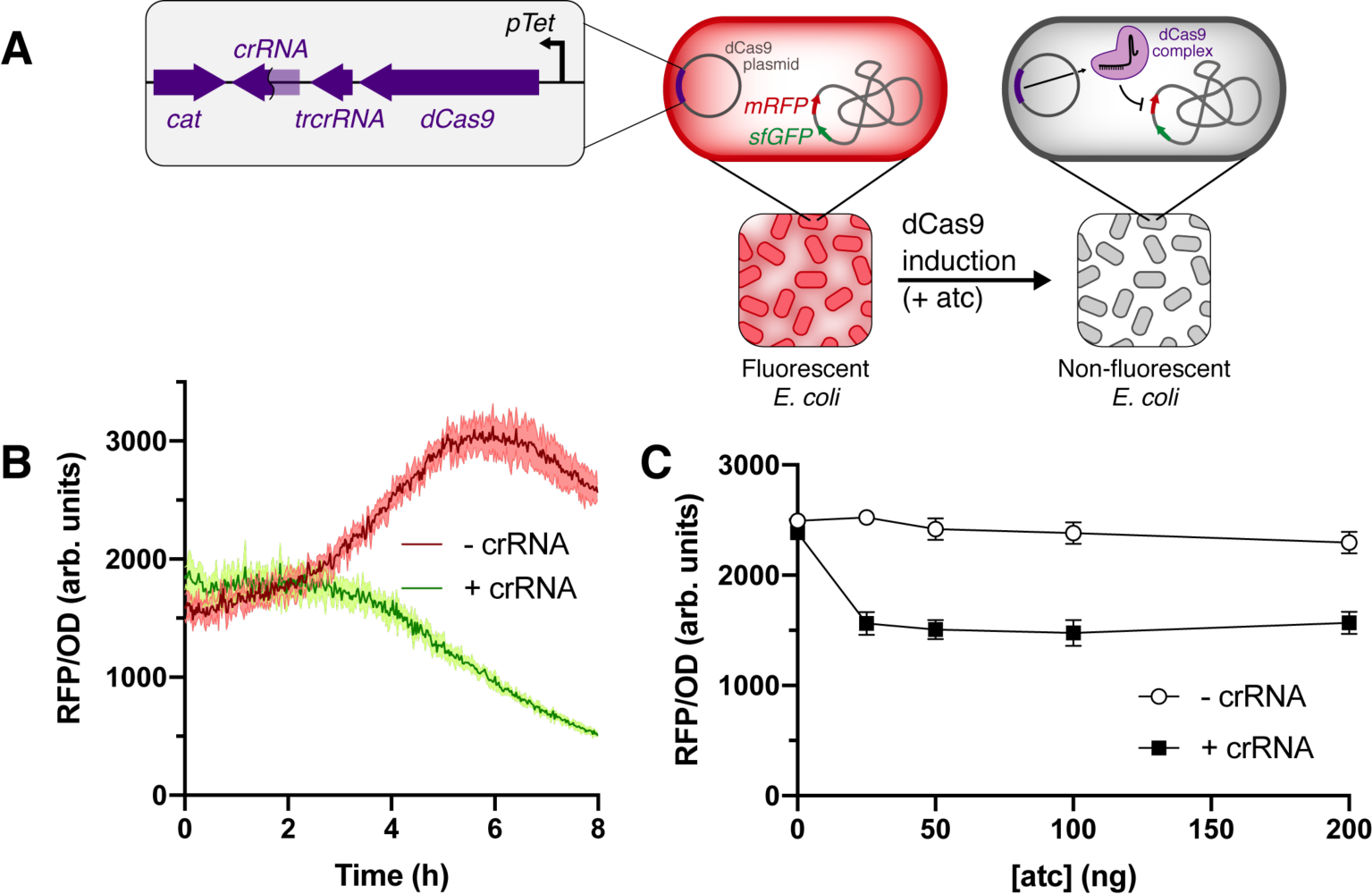
Plasmid based *in vitro* gene repression. *E. coli* transformed with atc inducible expression of dCas9 with and without crRNA targeting *rfp* (A). Relative fluorescence of over time after induction with 25 ng atc (B) and as a function of atc concentration after 6 h of incubation (C).

**Figure S3.**
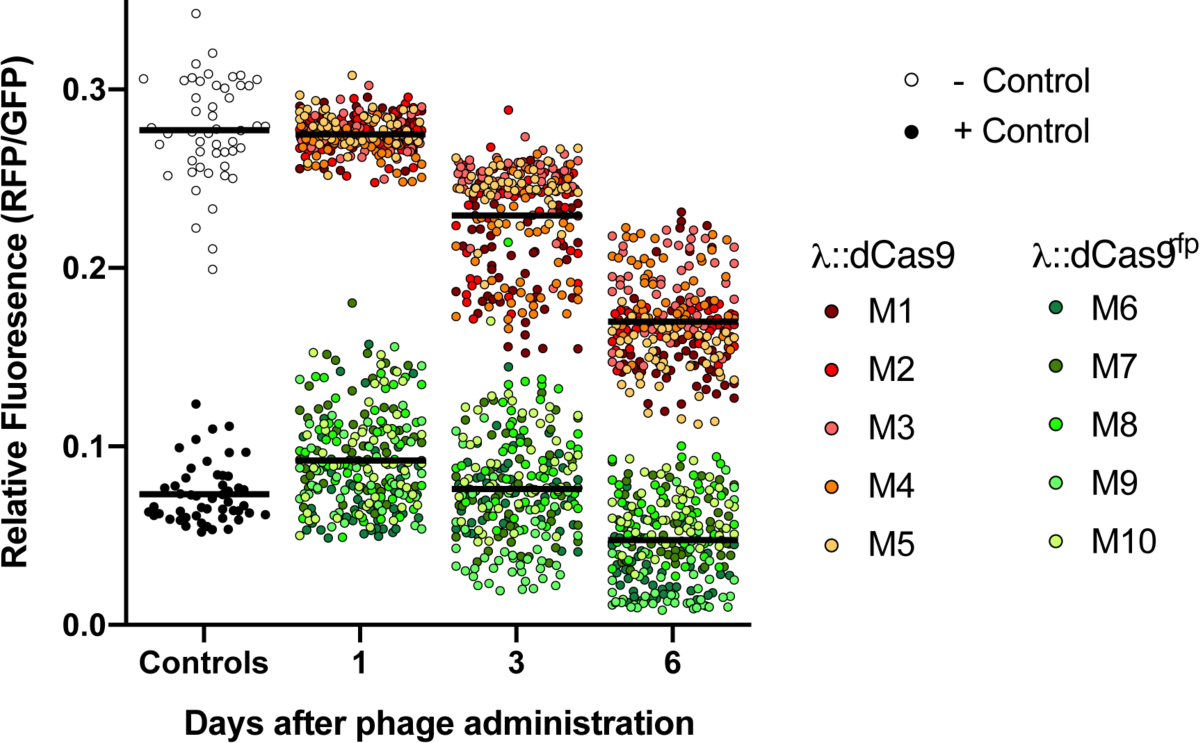
Ensemble view of relative colony fluorescence of fecal E. coli. 50 colonies from mice receiving either λ::dCas9 (n=5) or λ::dCas9^rfp^ phage (n=5). Controls represent lysogens after in vitro culture.

**Figure S4.**
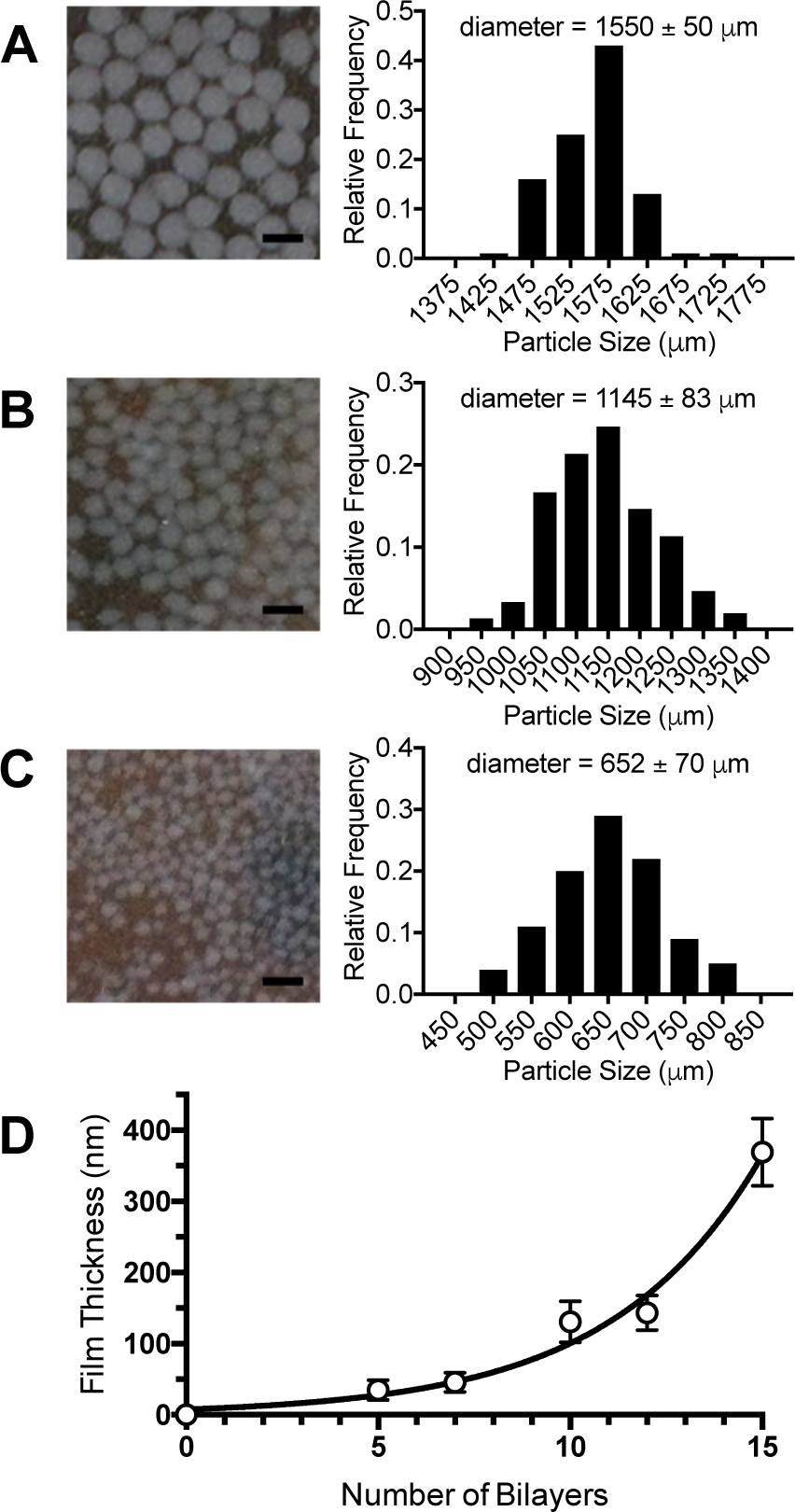
Properties of encapsulation formulation. Alginate beads are generated by dropwise addition of an alginate solution into an aqueous calcium chloride solution, assisted by coaxial air flow at 0 standard cubic feet per minute (scfm) (A), 50 scfm (B), and 100 scfm (C). The film thickness of the subsequent multilayer thin film of (polyethylenimine/pectin)_n_ was characterized on silicon wafers (D). Bead diameters and film thickness are represented as mean ± s.d.. Scale bars = 2 mm.

**Figure S5.**
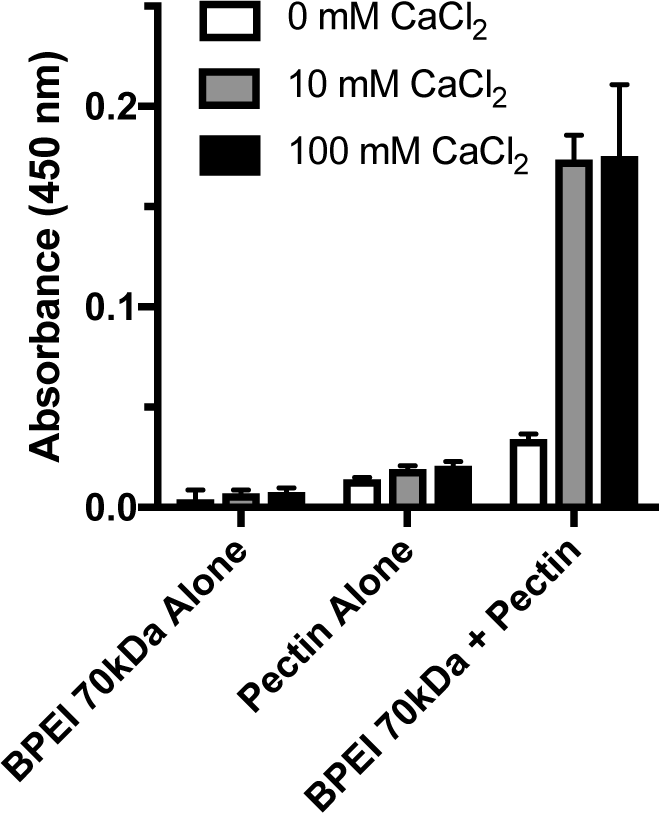
Polyelectrolyte complexation in calcium chloride. Absorbances at 450 nm of polyelectrolyte complexes between PEI and pectin in the presence of calcium chloride. Bars represent the mean ± s.d..

**Figure S6.**
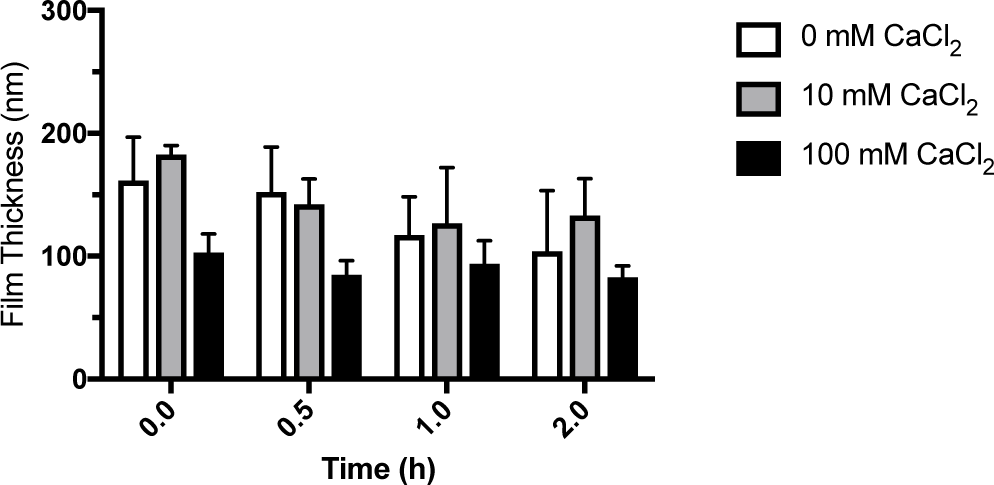
Thicknesses of films after incubation in SGF with pepsin. Films of (PEI/pectin)_10_ were assembled in 0 mM, 10 mM or 100 mM calcium chloride, then incubated in SGF, pH 1.1 with 1.5 mg/mL (5,000U/mL) pepsin at 37°C, and measured for thickness by profilometry. Bars represent the mean ± s.d..

**Figure S7.**
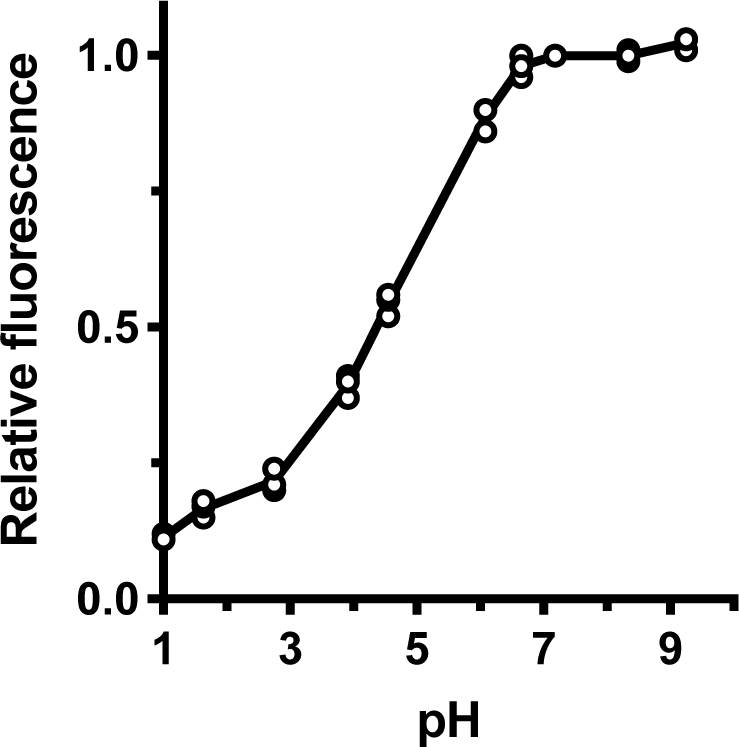
Fluorescence of Oregon Green 488-dextran. Relative fluorescence of the dye-polymer conjugate as a function of pH. Symbols represent replicates while the line represents the mean.

**Figure S8.**
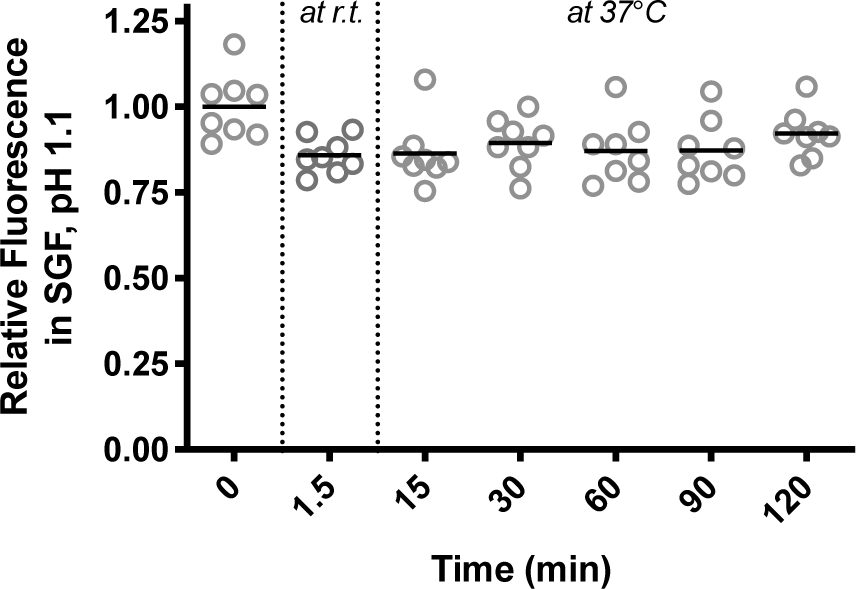
Change in fluorescence after prolonged incubation in SGF, pH 1.1. Alginate beads coated with (PEI/pectin)_10.5_ films were incubated for up to 2 hours at 37°C. The fluorescence from the encapsulated Oregon Green 488-dextran conjugate was measured periodically. Symbols represent individual beads and the bars indicate the means.

**Figure S9.**
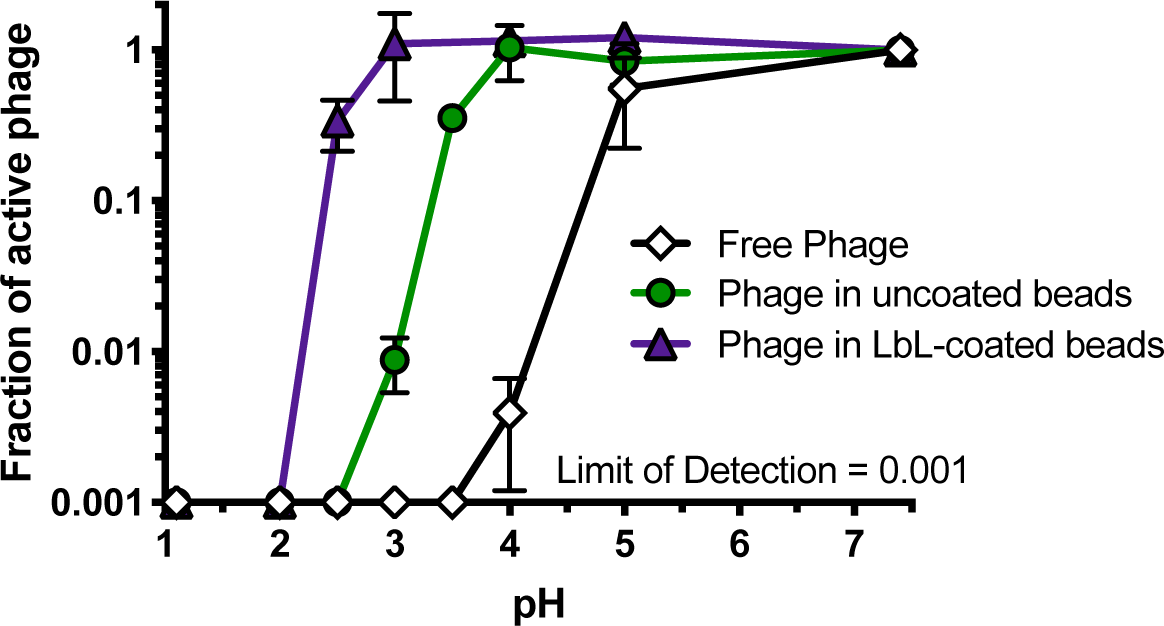
Bacteriophage stability at different pH conditions. Titration of bacteriophage free in solution, encapsulated in alginate beads, and encapsulated in alginate beads coated with (PEI/pectin)_15.5_ films incubated in SGF at various pH conditions. Data is represented as the mean ± s.d..

**Figure S10.**
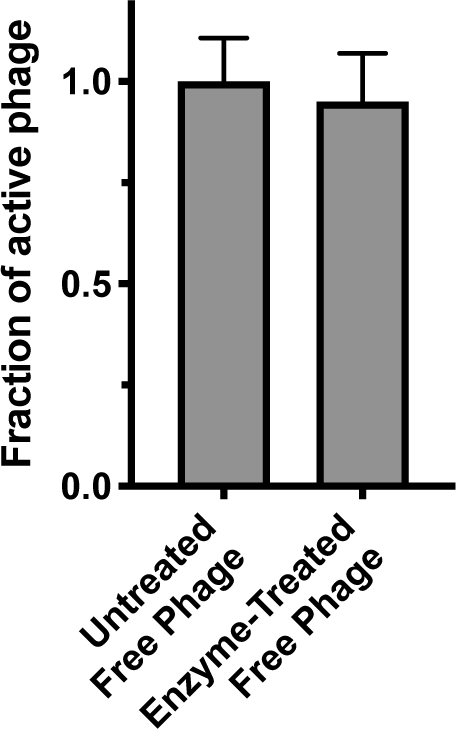
Treatment of bacteriophage with alginate lyase and pectin lyase. Influence of enzyme incubation on free phage. Bars represent the mean ± s.d..

**Figure S11.**
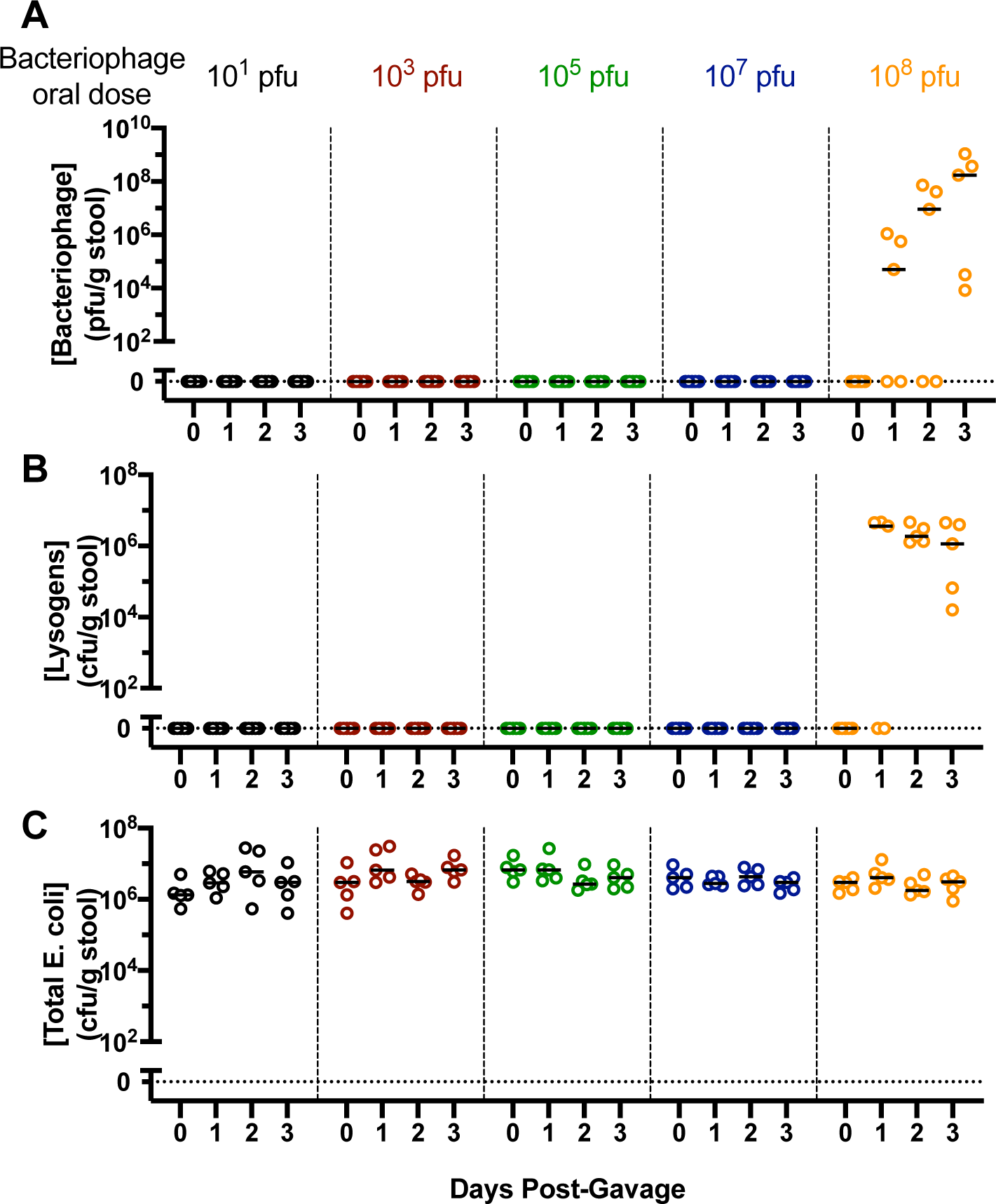
Inactivation of free bacteriophage administered orally to mice. After free bacteriophage suspended in water was administered by oral gavage to BALB/c mice, the stool was assayed for bacteriophage (A), lysogens (B), and total E. coli (C). Symbols represent individual mice (n = 5) and the lines represent the median.

**Figure S12.**
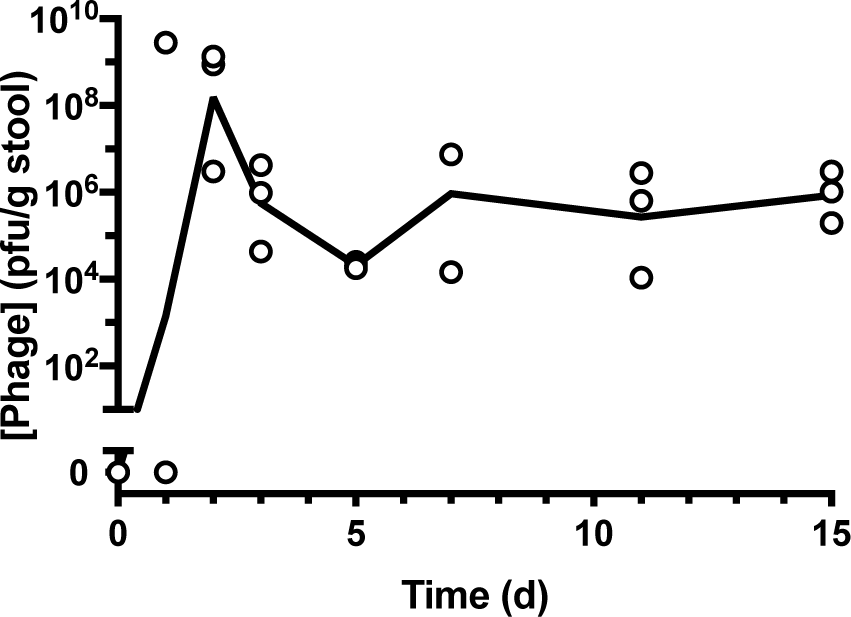
Duration of bacteriophage colonization in vivo. Bacteriophage encapsulated in (PEI/pectin)_15.5_ - coated alginate beads were orally administered to mice and show persistent colonization for at least two weeks. Symbols represent individual mice (n = 3) with the line indicating the medians.

**Figure S13.**
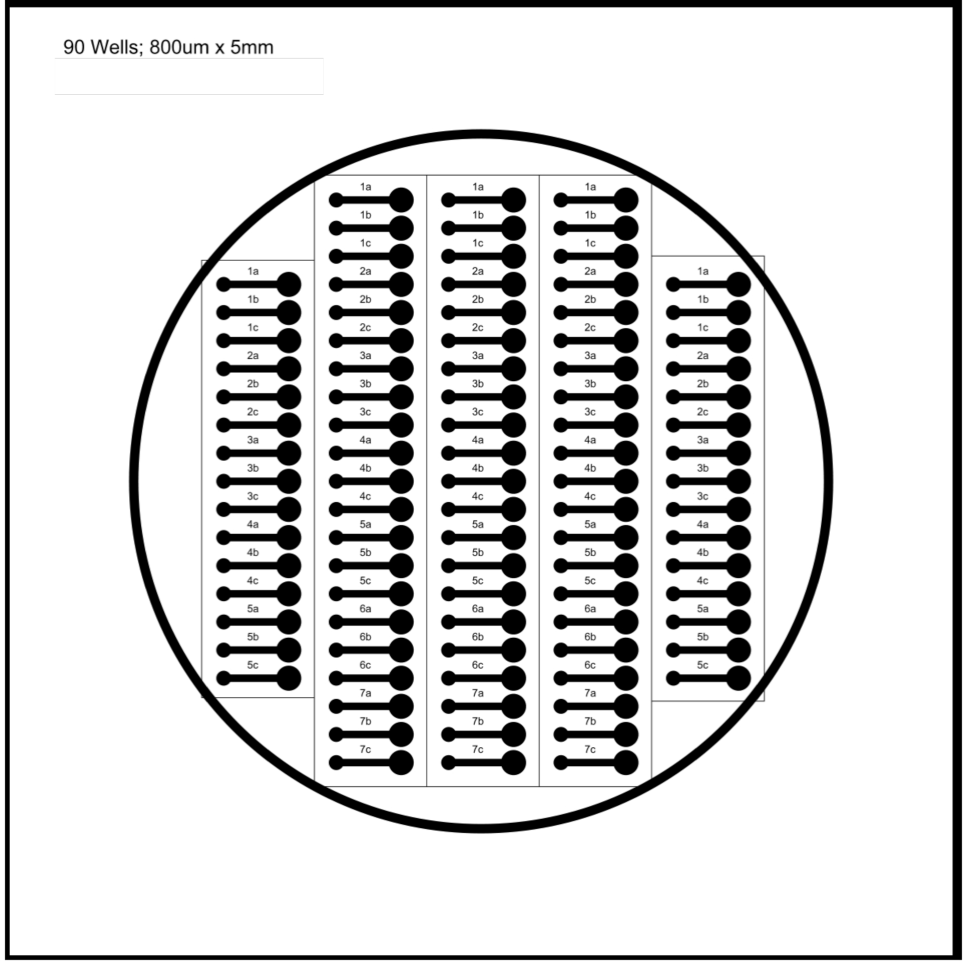
Photolithographic masks used for preparation of master templates.

## Bibliography

1. Gilbert, J. A. et al. Microbiome-wide association studies link dynamic microbial consortia to disease. Nature 535, 94–103 (2016).

2. Lynch, S. V. & Pedersen, O. The Human Intestinal Microbiome in Health and Disease. N. Engl. J. Med. 375, 2369–2379 (2016).

3. Zmora, N., Zeevi, D., Korem, T., Segal, E. & Elinav, E. Taking it Personally: Personalized Utilization of the Human Microbiome in Health and Disease. Cell Host Microbe 19, 12–20 (2016).

4. Smart, A. L., Gaisford, S. & Basit, A. W. Oral peptide and protein delivery: intestinal obstacles and commercial prospects. Expert Opin. Drug Deliv. 11, 1323–1335 (2014).

5. Layer, P., Jansen, J. B. M. J., Cherian, L., Lamers, C. B. H. W. & Goebell, H. Feedback Regulation of Human Pancreatic Secretion. Gastroenterology 98, 1311–1319 (1990).

6. Tennant, S. M. et al. Influence of Gastric Acid on Susceptibility to Infection with Ingested Bacterial Pathogens. Infect. Immun. 76, 639 (2008).

7. Imhann, F. et al. Proton pump inhibitors affect the gut microbiome. Gut 65, 740 (2016).

8. McConnell, E. L., Fadda, H. M. & Basit, A. W. Gut instincts: Explorations in intestinal physiology and drug delivery. Int. J. Pharm. 364, 213–226 (2008).

9. Hsu, B. B., Way, J. C. & Silver, P. A. Stable Neutralization of a Virulence Factor in Bacteria Using Temperate Phage in the Mammalian Gut. mSystems 5, e00013–20 (2020).

10. Dressman, J. B. et al. Upper Gastrointestinal (GI) pH in Young, Healthy Men and Women. Pharm. Res. 7, 756–761 (1990).

11. Evans, D. F. et al. Measurement of gastrointestinal pH profiles in normal ambulant human subjects. Gut 29, 1035 (1988).

12. Simon, G. L. & Gorbach, S. L. Intestinal flora in health and disease. Gastroenterology 86, 174–193 (1984). *45. Lambda II*. (Cold Spring Harbor Laboratory, 1983).

13. Friedland, A. E. et al. Characterization of Staphylococcus aureus Cas9: a smaller Cas9 for all-in-one adeno-associated virus delivery and paired nickase applications. Genome Biol. 16, 257 (2015).

14. Nishimasu, H. et al. Crystal Structure of Staphylococcus aureus Cas9. Cell 162, 1113–1126 (2015).

15. Qi, L. S. et al. Repurposing CRISPR as an RNA-Guided Platform for Sequence-Specific Control of Gene Expression. Cell 152, 1173–1183 (2013).

16. Lavalle, Ph. et al. Comparison of the Structure of Polyelectrolyte Multilayer Films Exhibiting a Linear and an Exponential Growth Regime: An in Situ Atomic Force Microscopy Study. Macromolecules 35, 4458–4465 (2002).

17. McConnell, E. L., Basit, A. W. & Murdan, S. Measurements of rat and mouse gastrointestinal pH, fluid and lymphoid tissue, and implications for in-vivo experiments. J. Pharm. Pharmacol. 60, 63–70 (2008).

18. Jończyk, E., Kłak, M., Międzybrodzki, R. & Górski, A. The influence of external factors on bacteriophages—review. Folia Microbiol. (Praha) 56, 191–200 (2011).

19. Cully, M. Microbiome therapeutics go small molecule. Nat. Rev. Drug Discov. 18, 569–572 (2019).

20. Wong, A. C. & Levy, M. New Approaches to Microbiome-Based Therapies. mSystems 4, e00122–19 (2019).

21. Reyes, A. et al. Viruses in the faecal microbiota of monozygotic twins and their mothers. Nature 466, 334–338 (2010).

22. Bondy-Denomy, J. & Davidson, A. R. When a virus is not a parasite: the beneficial effects of prophages on bacterial fitness. J. Microbiol. 52, 235–242 (2014).

23. Prüsse, U. et al. Comparison of different technologies for alginate beads production. Chem. Pap. 62, 364 (2008).

24. Castleberry, S. A., Li, W., Deng, D., Mayner, S. & Hammond, P. T. Capillary Flow Layer-by-Layer: A Microfluidic Platform for the High-Throughput Assembly and Screening of Nanolayered Film Libraries. ACS Nano 8, 6580–6589 (2014).

25. Chait, R., Shrestha, S., Shah, A. K., Michel, J.-B. & Kishony, R. A Differential Drug Screen for Compounds That Select Against Antibiotic Resistance. PLOS ONE 5, e15179 (2010).

